# Discovery of A Chimeric Polyketide Family as Cancer Immunogenic Chemotherapeutic Leads

**DOI:** 10.1101/2024.11.05.622009

**Authors:** Dan Xue, Mingming Xu, Michael D. Madden, Xiaoying Lian, Ethan A. Older, Conor Pulliam, Yvonne Hui, Zhuo Shang, Gourab Gupta, Manikanda K. Raja, Yuzhen Wang, Armando Sardi, Yaoling Long, Hexin Chen, Daping Fan, Tim S. Bugni, Traci L. Testerman, Qihao Wu, Jie Li

**Affiliations:** Department of Chemistry and Biochemistry, University of South Carolina, Columbia, South Carolina 29208, United States; Pharmaceutical Sciences Division, University of Wisconsin–Madison, Madison, Wisconsin 53705, United States; Department of Pathology, Microbiology and Immunology, School of Medicine, University of South Carolina, Columbia, South Carolina 29209, United States; Department of Biological Sciences, University of South Carolina, Columbia, South Carolina 29208, United States; Department of Cell Biology and Anatomy, School of Medicine, University of South Carolina, Columbia, South Carolina 29209, United States; Department of Surgical Oncology, The Institute for Cancer Care at Mercy, Mercy Medical Center, Baltimore, Maryland 21202, United States; Department of Biological and Physical Sciences, South Carolina State University, Orangeburg, SC 29117, United States; Lachman Institute for Pharmaceutical Development, University of Wisconsin–Madison, Madison, Wisconsin, United States

**Keywords:** natural products, activity metabolomics, MicroED, immunogenic chemotherapeutics, immunogenic cell death

## Abstract

Discovery of cancer immunogenic chemotherapeutics represents an emerging, highly promising direction for cancer treatment that uses a chemical drug to achieve the efficacy of both chemotherapy and immunotherapy. Herein we report a high-throughput screening platform and the subsequent discovery of a new class of cancer immunogenic chemotherapeutic leads. Our platform integrates informatics-based activity metabolomics for rapid identification of microbial natural products with both novel structures and potent activities. Additionally, we demonstrate the use of microcrystal electron diffraction (MicroED) for direct structure elucidation of the lead compounds from partially purified mixtures. Using this strategy to screen geographically and phylogenetically diverse microbial metabolites against pseudomyxoma peritonei, a rare and severe cancer, we discovered a new class of leads, aspercyclicins. The aspercyclicins feature an unprecedented tightly packed polycyclic polyketide scaffold that comprises continuous fused, bridged, and spiro rings. The biogenesis of aspercyclicins involves two distinct biosynthetic pathways, leading to formation of chimeric compounds that cannot be predicted by bottom-up approaches mining natural products biosynthetic genes. With comparable potency to some clinically used anticancer drugs, aspercyclicins are active against multiple cancer cell types by inducing immunogenic cell death (ICD), including the release of damage-associated molecular patterns and subsequent phagocytosis of cancer cells. The broad-spectrum ICD-inducing activity of aspercyclicins, combined with their low toxicity to normal cells, represents a new class of potential cancer immunogenic chemotherapeutics and particularly the first drug lead for pseudomyxoma peritonei treatment.

## Introduction

Conventional cancer chemotherapeutics are based on the cytotoxicity that also kills normal cells and are often immunosuppressive that prohibits immune responses from eliminating cancer cells, leading to side effects, drug resistance, and tumor regrowth^1,2^. On the other hand, cancer immunotherapy drugs stimulate host immunity to combat cancer but fail to induce immune responses in many patients^3,4^. Interestingly, it was recently discovered that some compounds can simultaneously exhibit cytotoxicity and induce immunogenicity against cancer, leading to immunogenic cell death (ICD) through activation of robust innate and adaptive anti-cancer immune responses^5,6^. Thus, discovery of immunogenic chemotherapeutics has emerged as an innovative direction for cancer treatment that combines the advantages of both chemotherapy and immunotherapy drugs^7,8^. Due to the intrinsic complexity of cancer immunology, rational design of immunogenic chemotherapeutics is currently immensely challenging. In contrast, a promising strategy is to discover natural immunogenic chemotherapeutics evolved by nature that have tremendous chemical diversities and intrinsic biological activities. This rationale is further strengthened by the well-known role of natural products in anti-cancer drug discovery and clinical use^9–17^, as well as the reported potential of natural products for inducing ICD^18^.

The key to effective screening of natural product libraries is both novelty and chemical diversity of the library. In order to build a chemically diverse library enriched in novel natural products, cutting edge cheminformatics tools, such as Global Natural Product Social Molecular Networking (GNPS)^19^ and Sum formula Identification by Ranking Isotope patterns Using mass Spectrometry (SIRIUS)^20,21^, have been increasingly used to mine unknown natural products and enable machine learning-based chemodiversity analyses. When combined with high-throughput screening (HTS), these tools can enable precise and effective mining of natural products with both novel structural features and targeted activities^22–24^, and thus guide targeted isolation of the compounds of interest. Another hurdle in natural product-based drug discovery is fast and unambiguous elucidation of complex chemical structures, particularly when confronted with limited material. This challenge can be addressed by the recent advances in MicroED, which enables the analysis of microcrystals even in a mixture, offering easier processing when compared to X-ray single crystal diffraction^25–28^.

To enhance screening efficiency and identify new immunogenic chemotherapeutic leads for a rare and sever cancer, pseudomyxoma peritonei (PMP) which currently lacks a drug discovery platform, we developed a PMP cell line along with an HTS platform. Our HTS platform (Figure S1) was integrated with cheminformatics and MicroED that facilitated the streamlined and efficient discovery of a new class of lead molecules, aspercyclicins (Figure 1A), with a putative new mechanism of action that appears to leverage both cytotoxicity and ICD. Aspercyclicins represent a chimeric scaffold combining two different biosynthetic pathways that cannot be predicted by bottom-up genomics approach and are also minor components of the extract that may be overlooked by conventional pipelines. Aspcercyclicins exhibited broad-spectrum activity against all four cancer cell lines used (Figure 1B), particularly against PMP cell line, with an activity comparable to clinically used anti-cancer drugs (Figure 1B). Aspercyclicins induced ICD, characterized by the release of immunostimulatory signals and the activation of antigen-presenting cells, representing the first immunogenic chemotherapeutic lead for PMP. In sum, the discovery of structurally unique and biologically active aspercyclicins underscores the value and potential of discovering promising cancer immunogenic chemotherapeutics from natural products chemical diversity. Importantly, this discovery also validates our innovative platform and integrated approach that can be widely applicable to the discovery of other biological activities of nature’s molecules.

**Figure 1.**
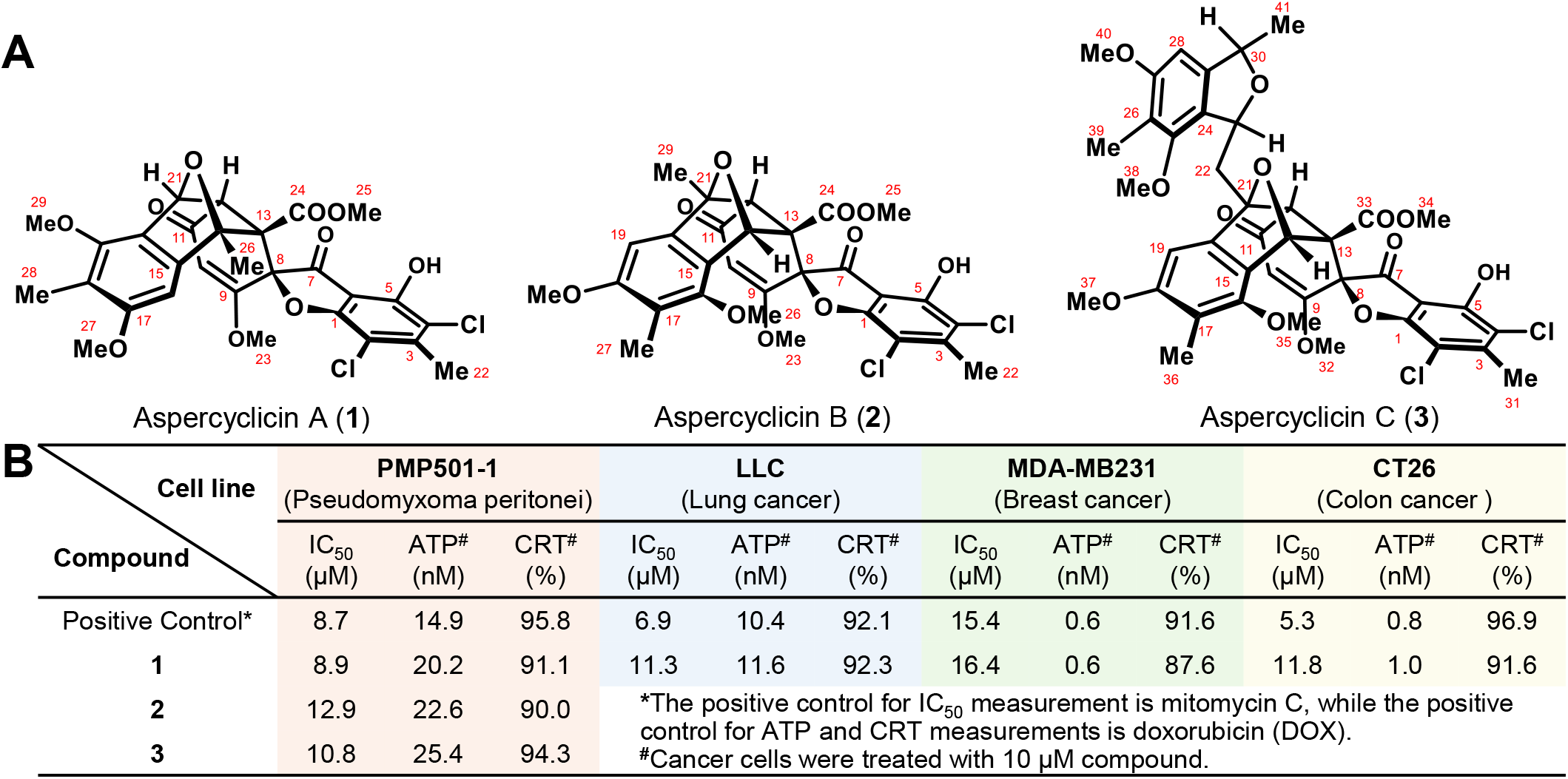
Aspercyclicins A (1), B (2) and D (3) isolated from *Aspergillus* sp. LI2020F001 simultaneously showed cytotoxicity and induced immunogenicity against cancer cell lines. (**A**) Chemical structure of aspercyclicins A-C (**1**-**3**). (**B**) The IC_50_ values of cytotoxicity and the induction of ICD hallmarks, including active secretion of adenosine triphosphate (ATP) and surface exposure of calreticulin (CRT), for aspercyclicins against four cell lines from both rare (pseudomyxoma peritonei) and common (lung, breast, and colon) cancers.

## Results

### Cancer Cell Line Development and Selection to Build A High-Throughput Platform for Immunogenic Chemotherapeutics Screening

Since there are no established PMP cell lines for drug screening, we first set out to establish a PMP cell line (Figure 2A) to support HTS and complement the commonly used cancer cell lines for testing (Figure 2B). We collected 56 primary PMP tumor samples from the peritonea of patients (Mercy Medical Center) with diagnosed PMP (Figure 2A and Figure S2). We successfully established cell lines from these peritoneal tumor tissues, marking the first development of PMP cell lines directly from the peritoneum. Previously, only three PMP-related cell lines were established, including N14A and N15A derived from appendiceal cancer tissue^29^ and NCC-PMP1-C1 originating from a distant PMP metastasis thigh tumor^30^. The N14A and N15A cell lines were immortalized with SV40, since they would not continue growing otherwise. Our cell lines were not genetically manipulated and represent a range of PMP subtypes. The appendix is the originating site of PMP; subsequently PMP disseminates widely in the abdomen but rarely metastasizes beyond the peritoneal cavity^31^. Considering this, N14A, N15A, and NCC-PMP1-C1 may not directly reflect the typical behavior of PMP. Therefore, it is imperative to establish PMP cell lines directly from PMP patient peritoneal tumor tissues to facilitate anti-PMP drug screening. Notably, three cell lines, PMP501-1 and PMP457-2, and ABX023-1, achieved successful passaging more than 50 times.

**Figure 2.**
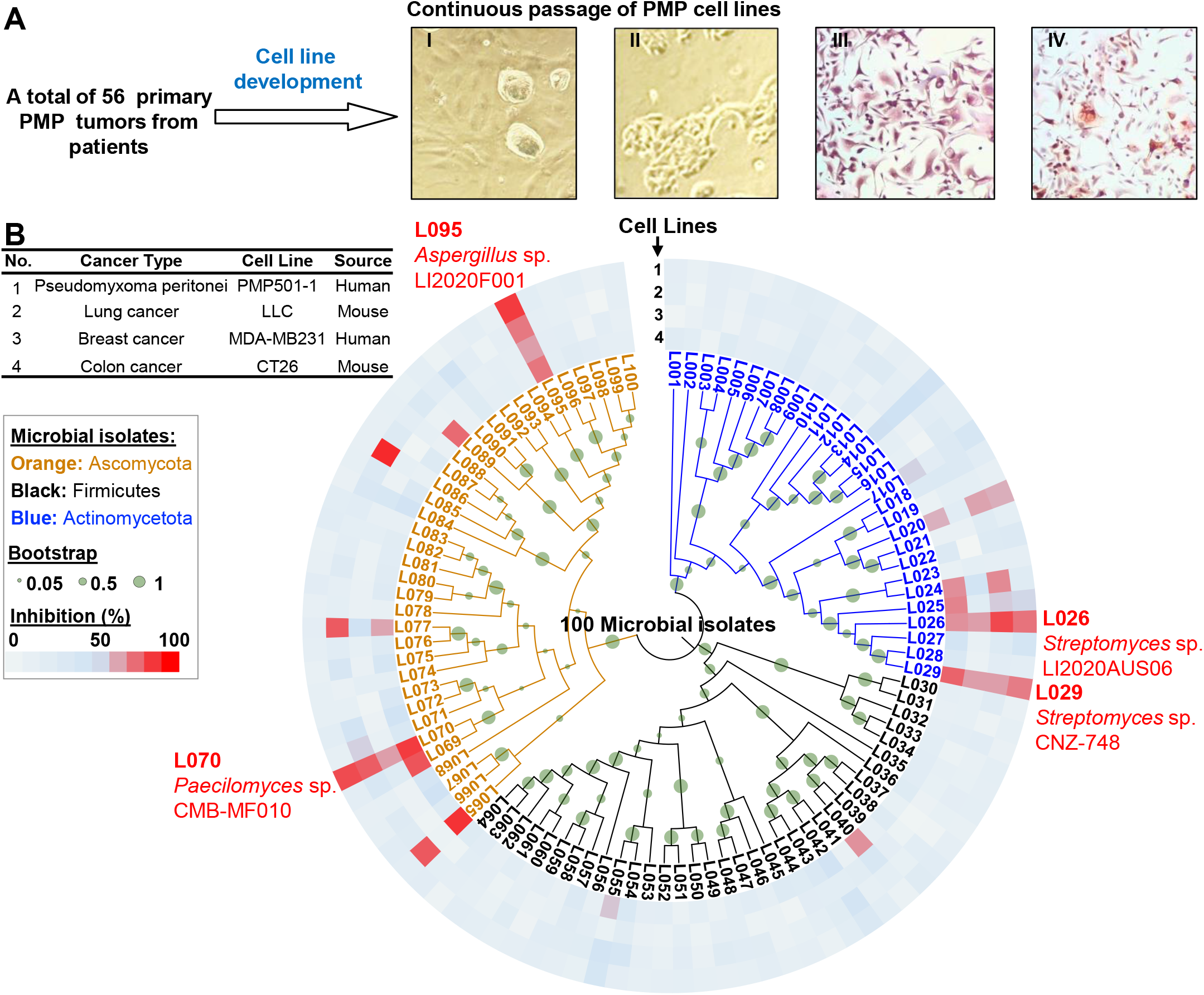
Cancer cell line development and selection followed by high-throughput activity screening of 100 microbial crude extracts. (**A**) Development of PMP cell lines for screening potential immunogenic chemotherapeutics. Continuous passage of PMP501-1 cell lines (as a representative example): I: Tumor cells with mostly mesenchymal phenotype at passage 6; II: The same cell line at passage 17 showing epithelial phenotype; III: MUC2 staining; IV: CEA staining. (**B**) A rapid, high-throughput cultivation and screening of microbial crude extracts against four cancer cell lines indicated the potential of microbial metabolites as anti-cancer leads. Four isolates, L026, L029, L070, and L095, were prioritized out of 100 strains tested because they caused strong cell death against all four cancer cell lines. The information of cancer cell lines is listed in the table.

Next, we established the best morphology of PMP cell lines for immunogenic chemotherapeutics screening. Epithelial cells exhibit the capacity to form monolayers, adhere effectively to culture surfaces, and thrive efficiently in defined media. These characteristics facilitate the implementation of high-throughput screening in a controlled environment, allowing for the comprehensive monitoring of cell behavior and responses to various compounds. Thus, we endeavored to establish stable PMP cell lines to facilitate further screening. In most cases, PMP tumor cells initially exhibit a mesenchymal morphology and require platelet derived growth factor (PDGF) to sustain their growth. Importantly, the initial cellular phenotype does not correlate with the specific tumor subtype, such as disseminated peritoneal adenomucinosis (DPAM) or peritoneal mucinous carcinomatosis (PMCA). Subsequently, cells undergo mesenchymal-epithelial transition (MET), changing from a mesenchymal to an epithelial morphology, starting from around passage six (Figure 2A). At this stage, the requirement for PDGF diminishes, but the necessity for insulin remains unchanged. It is worth noting that we have not observed any instances of cell lines reverting from an epithelial to a mesenchymal state, even after undergoing more than fifty passages. Three cell lines (PMP501-1, PMP457-2, and ABX023-1) with successful passaging also showed stable epithelial morphology. Two of them (PMP501-1 and PMP457-2) were from patients with highly differentiated DPAM tumors and one (ABX023-1) was a highly differentiated tumor with signet ring morphology, representing the form of PMP with the worst prognosis. Thus, we selected PMP501-1, along with one lung cancer (LLC), one breast cancer (MDA-MB231), and one colon cancer (CT26) cell lines, for subsequent bioactivity screening.

### Screening of A Strain Library Encompassing a Broad Spectrum of Geographic and Phylogenetic Diversity

With the selected and newly established cancer cell lines, we assembled a microbial library containing 100 bacterial and fungal isolates. The isolates were selected based on several features including geographical and phylogenetic diversity (Supplementary File S1): i). Representing both an in-house collection (56 strains) and commercial sources (44 strains); ii). Originating from a wide range of geographic location (Asia, North America, South America, Europe, and Australia); iii). Including 36 fungal and 64 bacterial strains renowned for their prolific production of natural products; and iv). 61 of these strains being of marine or human microbial origin, both of which are considered underexplored sources of biologically active natural products. For the isolates from the in-house collection, we identified the strains based on the 16S rRNA sequences (bacteria) or ITS regions (fungi) (Supplementary File S1). We set out to first screen cytotoxicity of the extracts of this microbial library against the cancer cell lines selected. By combining phylogenetic analysis and cytotoxicity assay results (Figure 2B, Figure S3, and Table S1), we observed intriguing patterns that guided further screening efforts, including: i). Extracts of phylogenetically distinct strains could exhibit comparably strong bioactivity, suggesting the potential of unearthing microbial metabolites to combat cancer; ii). Approximately half of the tested *Streptomyces* strains displayed a cell inhibition efficacy of 50% or greater (Figure 2B) against at least one cell line, which is consistent with this genus being recognized as a prolific producer of bioactive metabolites; and iii). Strains from marine environments, both bacterial and fungal (*e*.*g*., L026, L029, L070, and L095 highlighted in red in Figure 2B), appeared to produce metabolites with better activity than those from other sources. Thus, we prioritized four strains that exhibited over 50% cytotoxicity at 10 μg/mL against all four selected cancer cell lines (Figure S3, Table S1), including L026 (*Streptomyces* sp. LI2020S06), L029 (*Streptomyces* sp. CNZ-748), L070 (*Paecilomyces* sp. CMB-MF010), and L095 (*Aspergillus* sp. LI2020F001), for subsequent activity metabolomics analysis to identify the compounds responsible for cancer cell death.

### Bioactivity Metabolomics Led to the Discovery of Aspercyclicins, A New Class of Minor Components with Strong Activity Correlation

We fractionated the crude extracts of each prioritized strain into four fractions by reversed-phase liquid chromatography. To directly target active compounds in fractionated samples, we conducted bioactivity-based molecular networking^32^ using cell death-based assays (Figure 3A, Figure S4-S7, and Tables S2). Bioactivity scores for all molecular features (MOFs) were calculated using their relative abundances along with each fraction’s corresponding bioactivity against cancer cells. Through this approach, 34 MOFs with statistically significant bioactivity scores [strong Pearson correlation coefficient (*r* > 0.99, largest nodes) and significance (*p* value < 0.001)] were selected from these fractions for further analysis (Table S3). Among them, major components exhibiting high bioactivity scores (*r* > 0.99, *p* value < 0.001) and annotated as known compounds, including actinomycins, grincamycins, and viridicatumtoxins, were identified from strains L026, L029, and L070, respectively (Figure 3A and Figure S8). These compounds have been previously reported with cytotoxic activity^17,33,34^, confirming the effectiveness of our bioactivity-based molecular networking analysis. Furthermore, we screened all cytotoxic fractions for ICD-inducing activity through measuring the hallmarks of ICD^5,6,35^ such as the active secretion of adenosine triphosphate (ATP) and the surface-exposure of calreticulin (CRT). The active fraction (Fr. C) of L095 (*Aspergillus* sp. LI2020F001) not only induced robust cell death (Figure 3A) but also exhibited potent ICD effects (Figures 3B and 3C). Specifically, it significantly increased extracellular ATP levels in PMP501-1 and LLC cell lines (p<0.0001) as well as the exposure of CRT on the surface of all four tested cell lines. These results motivated us to characterize the active compounds present in this fraction as potential leads for inhibiting PMP by inducing ICD.

**Figure 3.**
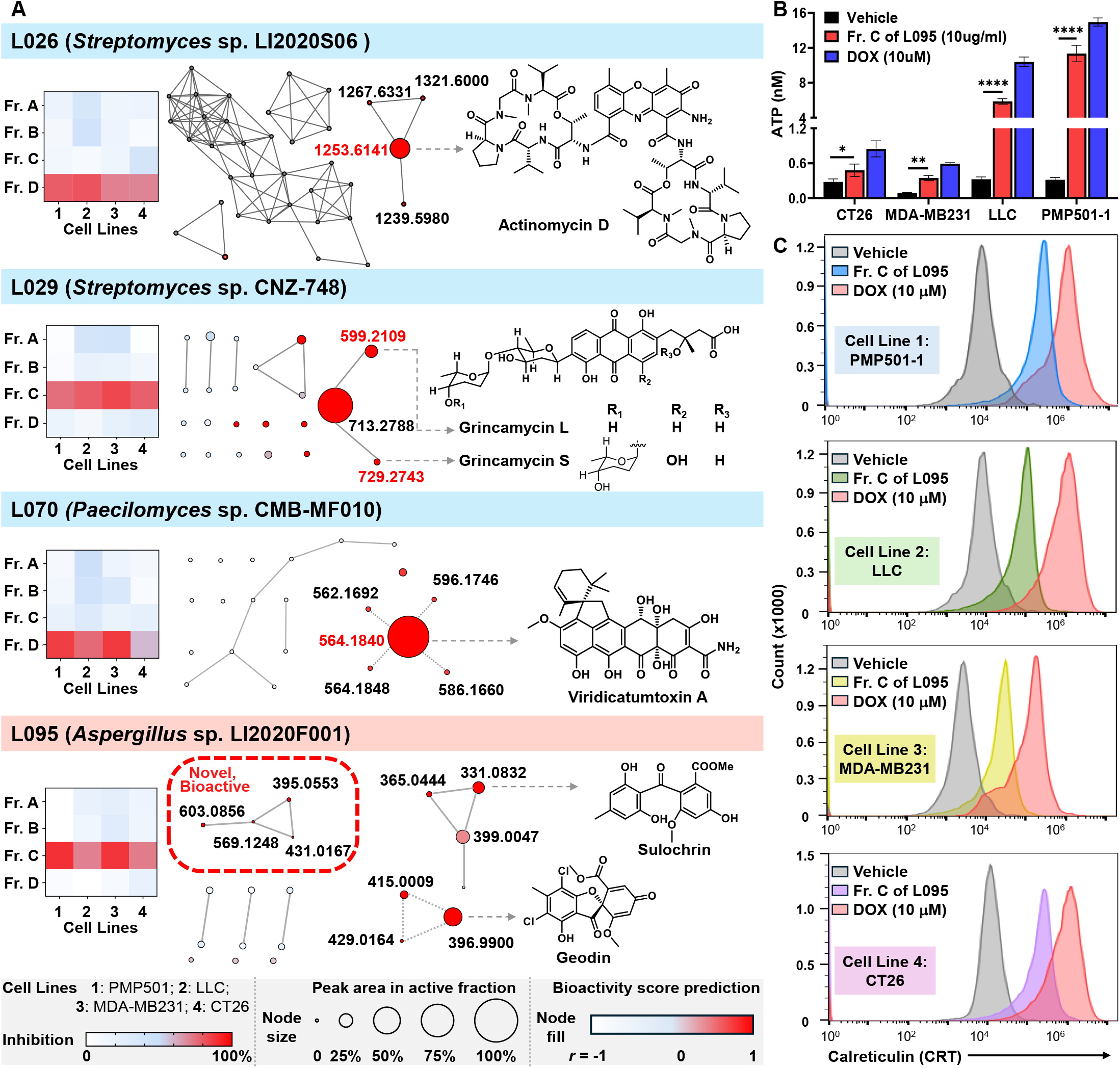
Bioactivity-based molecular networking reveals chemical features from strain L095 inhibiting cancer cell lines through induced immunogenicity. (**A**) Activity profiling of fractions from strain L026, L029, L070, and L095 against four cancer cell lines. Known active compounds were annotated through GNPS and MS2 analysis (Figure S8). In contrast, undescribed compounds with high bioactivity score from strain L095 were prioritized and highlighted in a red box. (**B**) The secretion of ATP of active fraction Fr. C of L095 against all four cancer cell lines indicated a possible anti-cancer effect through induced immunogenicity. (**C**) CRT measurement exhibited the potent ICD effect of Fr. C of L095 against four cancer cell lines.

After inspecting the molecular network of Fr. C of strain L095, only geodin, a previously reported *Aspergillus* metabolite^36^ and its analogs, including sulochrin, (Figure 3A) were annotated. However, geodin and its analogs are not known as potent anticancer agents. In fact, geodin only showed weak inhibitory activity (26.5% inhibition at a concentration of 10 μM) against PMP 501-1 in our bioassay (Figure S9). Thus, it suggested that Fr. C contained other unidentified minor components that contributed significantly to the observed cytotoxicity and ICD-inducing activity. While searching for the metabolite(s) contributing to the bioactivity, multiple MOFs with the same ion at *m/z* 603.0854 ([M - H]^−^) in Fr. C, which we termed aspercyclicins, caught our attention as we performed MS-based, SIRIUS-assisted analysis. Particularly, we observed that all aspercyclicins shared a tandem MS product ion at *m/z* 396.9897 that was directly assigned as geodin using SIRIUS (*m/z* 396.9898, [M - H]^−^, 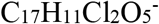, calc’d *m/z* 396.9887). This prediction was substantiated by similar MS/MS fragments between aspercyclicins and geodin (Figure S10). As such, we hypothesized that aspercyclicins were complex conjugates of geodin and an undescribed structural motif (C_12_H_14_O_3_, Figure S11). More importantly, our bioactivity-based molecular networking analysis revealed that the presence of aspercyclicins showed a strong correlation with the observed activity (Figure 3A).

### MicroED Directly Elucidated the Chemical Structure of Aspercyclicin A from A Highly Bioactive Fraction

Upon further fractionation of the active fraction Fr. C, a subfraction named Fr. C_sub was obtained which contained aspercyclicins and maintained the bioactivity (Figure S12). We next employed MicroED as a powerful tool^25,28,37,38^ to analyze this semi-pure mixture, with the goal of directly unraveling the structure of compounds in the active samples. A significant advantage of MicroED lies in its capability to analyze small crystals that may be unsuitable for conventional X-ray methods. Moreover, it has been demonstrated that even minuscule quantities of a natural product isolate^27^ or mixtures^25^ are sufficient for achieving comprehensive structural determination. Such an approach can effectively prioritize subsequent isolation efforts and expedite the structure elucidation of challenging compounds. We grew the microcrystals of the semi-pure active fraction through a gradual evaporation process. MicroED elucidated the full structure of two known metabolites, geodin and sulochrin (Figure 4, CCDC 2368919 and CCDC 2368920, Figures S13-S21, and Tables S4-S10) in the active fractions, confirming our metabolomics analysis results. More importantly, MicroED analysis revealed the full structure of one of the aspercyclicins, named aspercyclicin A (**1**, Figure 4, CCDC 2323844, Figures S22-S25, and Table S11-S13). The analysis confirmed the presence of geodin, *O*-methyl, and methyl groups, thereby corroborating our earlier MS-based cheminformatics analyses. Based on the structural analysis from MicroED, aspercyclicin A (**1**) contains a crowded 6/5/5/6/5/6 ring system that originates from the conjugation of a geodin and an unusual 1,4-dihydro-1,4-epoxynaphthalene motif. Aspercyclicin A (**1**), with its densely arranged polycyclic framework incorporating an oxygen-bridged ring and a spiro ring, represents the first example of complex natural products with an unprecedented hexacyclic skeleton. Motivated by this structural novelty revealed by MicroED and aspercyclicins’ correlation with activity demonstrated by bioactivity-based molecular networking, we purified aspercyclicin A (**1**) (Figure S26) and confirmed its cytotoxicity against PMP501-1 (IC_50_ 8.9 μM), comparable to the clinical PMP chemotherapeutic drug mitomycin C (IC_50_ 8.7 μM) (Figure 1), which in turn motivated us to identify more congeners of aspercyclicin A (**1**).

**Figure 4.**
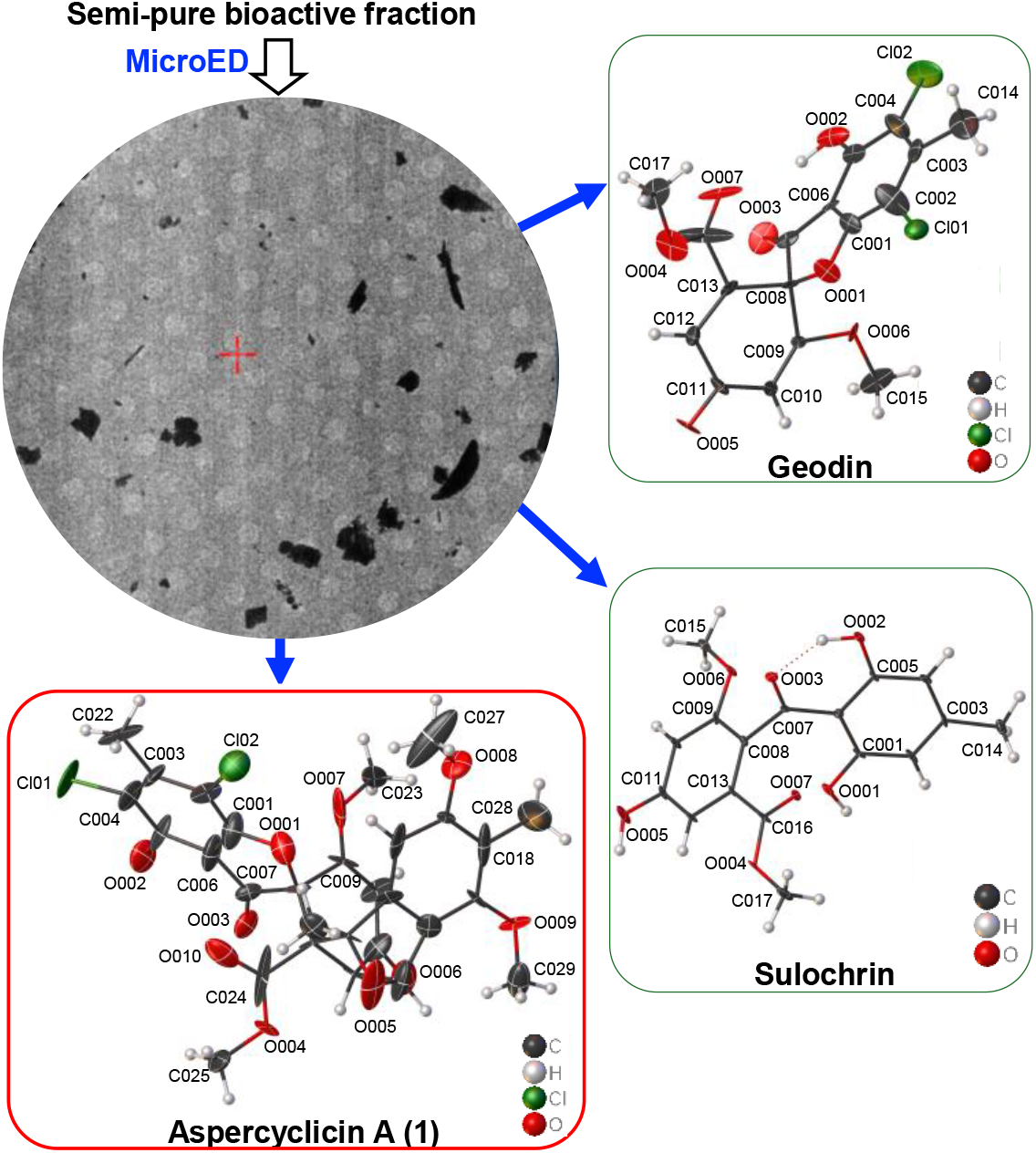
Direct structural elucidation of compounds in mixture from the active fraction Fr. C of strain L095 (*Aspergillus* sp. LI2020F001) by MicroED. The semi-pure active fraction was subjected to MicroED analysis. Crystal structures of geodin, sulochrin, and aspercyclicin A (**1**), determined by MicroED experiments, are depicted. The structures were solved to 0.80 Å, 0.65 Å, and 0.90 Å resolution for geodin, sulochrin, and aspercyclicin A (**1**), respectively.

### Chemical Diversity Expansion and Chimeric Biogenesis of Aspercyclicin Family

We conducted an extensive metabolomics analysis^39^, utilizing molecular networking and SIRIUS (Figure 5A and Figure S27), to identify aspercyclicin A (**1**) analogs and increase the chemical diversity of aspercyclicin family. Besides, inspired by building blocks-based molecular network strategy, we also took a typical MS^2^ geodin fragment of aspercyclicins (*m/z* 396.9897) as a probe to manually search for aspercyclicin analogs (Figure S11). As a result, we cheminformatically deduced the chemical structure of seven analogs of aspercyclicin A (**1**) (Figure 5B). Among them, MOF **2**, designated as aspercyclicins B, displayed the same molecular formula and a similar retention time, leading to the prediction that it might be a regio- or stereoisomer of aspercyclicin A (**1**). MOF **3**, designated as aspercyclicin C (**3**), is a conjugate of aspercyclicin A (**1**) with unidentified structural motifs. Other aspercyclicins, such as MOFs **4**–**7**, are likely shunt products with less choline atoms on the geodin residue. These novel features of these analogs motivated our isolation of these aspercyclicin analogs (Figure S26). We report here the NMR assignment of compound **1** (Figure 5C, Figures S28-S36, Table S14, and Supporting Information Chemical Structure Elucidation) and the *de novo* structure determination of **2**–**4** (Figure 5C, Figures S37-S55, Tables S14 and S15, and Supporting Information Chemical Structure Elucidation). The ^1^H and ^13^C NMR data of compound **2** showed great similarity to those of **1**, which indicated that they shared a similar polycyclic structure. However, compared to **1**, compound **2** exhibited a different substitution pattern of methyl groups on part B (Figure 5C). Compounds **3** and **4** exhibited an additional set of NMR signals that could be assigned as an additional 5,7-dimethoxy-1,6-dimethyl-1,3-dihydroisobenzofuran motif, which is attached to aspercyclicin B (**2**) through a C-C bond (Figures S44-S55). Compound **2** was also analyzed by the X-ray single crystal diffraction (Figure 5D, CCDC 2227218; Figures S56 and S57, and Tables S16-S23), which not only validated earlier structural hypotheses based on NMR spectroscopy but also revealed the absolute configuration of **2**. Due to limited material for X-ray or microED analysis, we tentatively assigned the stereochemistry of other aspercyclicins as depicted (Figure 5B) through 2D NMR and biogenetic logics. Collectively, the aspercyclicins discovered represent three different types of novel skeletons (Figure 5B).

**Figure 5.**
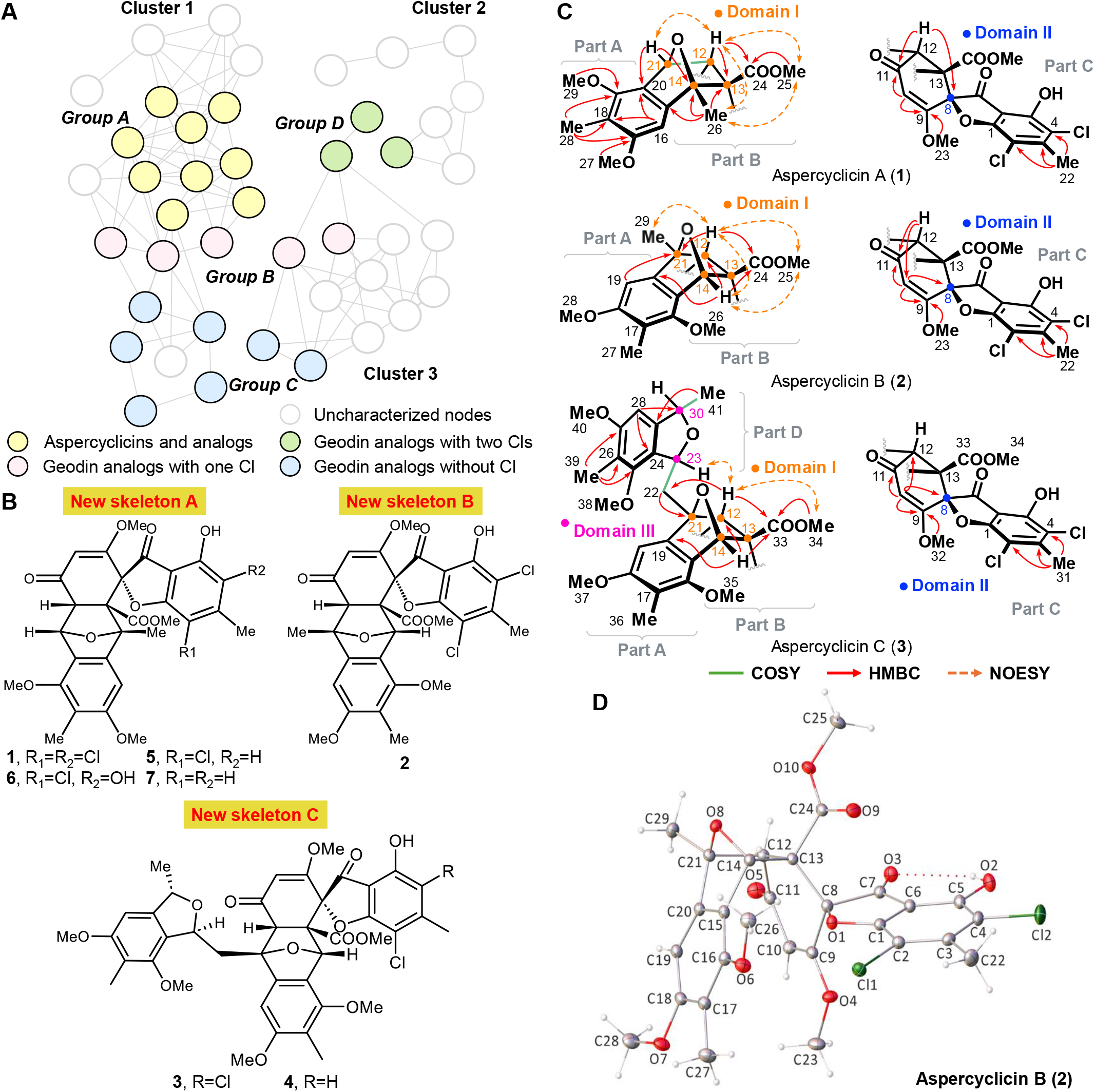
GNPS-based aspercyclicin analog discovery and structural elucidation. (**A**) A selected view of the molecular networking analysis of metabolites in strain L095. Molecular networking reveals three distinct clusters (clusters1-3), indicating the presence of structurally diverse analogs of aspercyclicin A (**1**). The identified MOFs were further divided into four groups and color-coded based on their chemical properties. MOFs in group A within cluster 1, highlighted in yellow, were identified as analogs of aspercyclicin A (**1**). MOFs in pink (group B) were analogs of geodin with one Cl atom, in blue (group C) were geodin analogs without Cl, and MOFs in green (group D) were geodin analogs with two Cl atoms. (**B**) Cheminformatically deduced chemical structures of aspercyclicin analogs. (**C**) NMR assignment of aspercyclicin A (**1**, top), aspercyclicin B (**2**, middle), and aspercyclicin C (**3**, bottom). ^1^H–^1^H COSY, key HMBC, and NOESY correlations for **1** (top), **2** (middle), and **3** (bottom). (**D**) X-ray crystal structure of aspercyclicin B (**2**), illustrating its absolute configuration.

Aspercyclicins A and B (**1** and **2**) feature two structural domains, comprising a geodin moiety linked to a 1,4-dihydro-1,4-epoxynaphthalene motif through C-C bonds and a bridged oxygen. Aspercyclicins C and D (**3** and **4**) contains an additional structural domain of 5,7-dimethoxy-1,6-dimethyl-1,3-dihydroisobenzofuran through a C-C linkage. We anticipate that the formation of these compounds results from the coupling of independent structural motifs, thus suggesting that their structures could be chimeric products of different biosynthetic gene clusters (BGCs). We sequenced the genome of *Aspergillus* sp. LI2020F001 and examined its BGCs with antiSMASH 7.0^40^. As expected, a type I polyketide synthase (T1PKS) pathway, named BGC1, exhibited significant similarity to the genes associated with the biosynthesis of geodin^41^ (Figure 6A), which presumably accounts for the generation of the geodin motif in aspercyclicins. In addition, the presence of a highly oxygenated phenethyl motif in aspercyclicins is reminiscent of ustethylin A^42^ and marilone A^43^, both of which are polyketides derived from *Aspergillus ustus* 3.3904 and *Stachylidium* sp. 293K04, respectively. The differences between the oxygenated phenethyl motif in aspercyclicins and ustethylin A were the *O*-methylation and oxidation positions. We further identified another T1PKS pathway, named BGC2, that has not been connected to any known natural products but shares partial similarity to the gene cluster (*utt*) encoding ustethylin A (Figure 6A), suggesting its potential role in producing the observed highly oxygenated phenethyl motif. We propose that the common intermediate ustethylinic acid is formed first (Figure 6B), which is reduced to an aldehyde product (I) by the NRPS-like gene of BGC2, analogous to the biosynthesis of ustethylin A^42^. The aldehyde (I) then forms a hemiacetal intermediate (II) followed by dehydration to form a furan intermediate (III) ^44–47^. Intermediate III could then engage in either unknown enzymatic or spontaneous coupling with the trisubstituted double bond in geodin through a Diels-Alder reaction, generating aspercyclicins A and B (**1** and **2**) that represent two types of novel skeletons^48^ (Figure 6B). Moreover, we hypothesize that a double bond rearranged intermediate (IV, Figure 6B), derived from intermediate III, can react with the aldehyde intermediate (I) *via* Michael addition, generating a conjugated diisobenzofuran intermediate V (Figure 6B and Figure S58). The following Diels-Alder coupling between intermediate V and geodin results in the formation of aspercyclicin C (**3**) that represents another type of novel skeleton (skeleton C). The producing strain of aspercyclicins employs the “crosstalk” between two distinct BGCs to assemble basic building blocks and couple them into chimeric natural products through novel machineries, thereby enhancing both structure and pharmacophore diversity. These chimeric products would elude bottom-up genome mining efforts and their targeted discovery showcases the power of our screening strategy. Further in-depth investigation is needed to verify the hypothesized biosynthesis leading to the formation of these new pharmacophores that inhibit PMP cancer cells.

**Figure 6.**
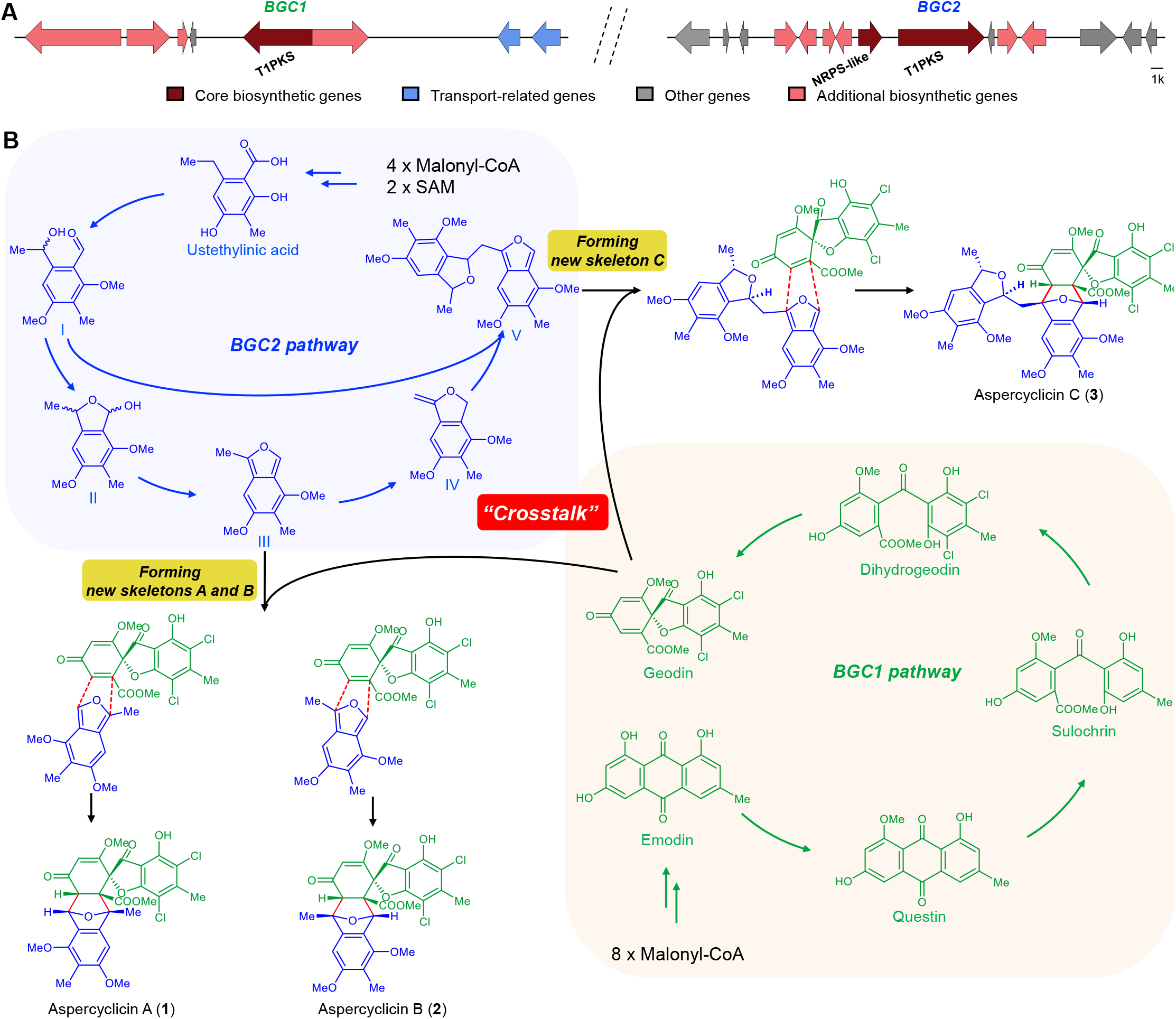
Proposed biosynthetic pathway for aspercyclicins. (**A**) Overview of the two gene clusters associated with the biosynthesis of aspercyclicins in L095 (*Aspergillus* sp. LI2020F001). Tentative functional annotations for genes and their relative organization are indicated by color as shown. (**B**) The proposed biosynthetic pathway of aspercyclicins reveals a “crosstalk” between two distinct polyketide pathways, resulting in the formation of four novel chimeric polyketide skeletons.

### Aspercyclicins Leveraged Both Cytotoxicity and ICD against Cancer Cells

At a concentration of 20 μM, aspercyclicins A–C (**1**–**3**) induced over 75% cell death across all tested cell lines (PMP501-1, LLC, MDA-MB231, and CT26; Figure S59 and Figure S60). Notably, they exhibited the highest cytotoxicity against PMP501-1, with IC_50_ values of 8.9, 12.9 and 10.8 μM, respectively (Figure 1B). To compare aspercyclicins with other therapeutic agents, we included diprovocim-1, paclitaxel, mitomycin C, and doxorubicin in our cytotoxicity assay. Both diprovocim-1 and paclitaxel demonstrated notably lower cell inhibition activity against PMP501-1 compared to aspercyclicin A (**1**) at both 10 and 20 μM, respectively (Figure S59). Mitomycin C, doxorubicin, and aspercyclicin A (**1**) exhibited comparable cytotoxicity against all tested cancer cell lines, but aspercyclicin A (**1**) showed much lower cytotoxicity towards normal cells compared to mitomycin C and doxorubicin, including human umbilical vein endothelial cells (HUVECs), H9c2 cells derived from embryonic rat cardiomyocytes, and the monocyte/macrophage-like cell line Raw264.7 (Figure S59).

We further explored the potential of aspercyclicins as immunogenic chemotherapeutics. Since aspercyclicin A (**1**) showed the most potent cytotoxic activity and weak cytotoxic activity to normal cells, we used **1** as a representative for study. We first explored whether **1** can induce the hallmarks of ICD, including the active secretion of ATP and CRT. Our results revealed that **1** induced an increase of ATP secretion in PMP501-1 and LLC cells (Figures 1B and 7A) as well as CRT exposure in all the four cell lines (Figures 1B and 7B), with the strongest activity observed in PMP501-1. Thus, we also tested compounds **2** and **3** in PMP501-1, with similarly active induction of ATP and CRT observed (Figure 1B). Together, these results suggested that aspercyclicin-induced cell death might be associated with ICD. We next explored potential signaling pathways involved in the observed induction of ICD. Previous studies have demonstrated that the exposure of CRT on the plasma membrane necessitates the induction of endoplasmic reticulum (ER) stress^49^. ER stress can be initiated by transmembrane sensor protein kinase RNA-like endoplasmic reticulum kinase (PERK)^50^, which further activates downstream signaling pathway. We observed that the expression of total PERK and its substrate eukaryotic translation initiation factor 2α (eIF2α) in PMP501-1 cells were not affected by treatment with **1** (Figure 7C). However, compound **1** enhanced phosphorylation levels of PERK and eIF2α in PMP501-1 cells. In addition, the expressions of C/EBP homologous protein (CHOP) downstream of the signaling pathway were increased by treatment of **1** (Figure 7C). Taken together, aspercyclicin-induced ICD might be mediated through the PERK/elF2α/ATF4/CHOP pathway (Figure S61).

**Figure 7.**
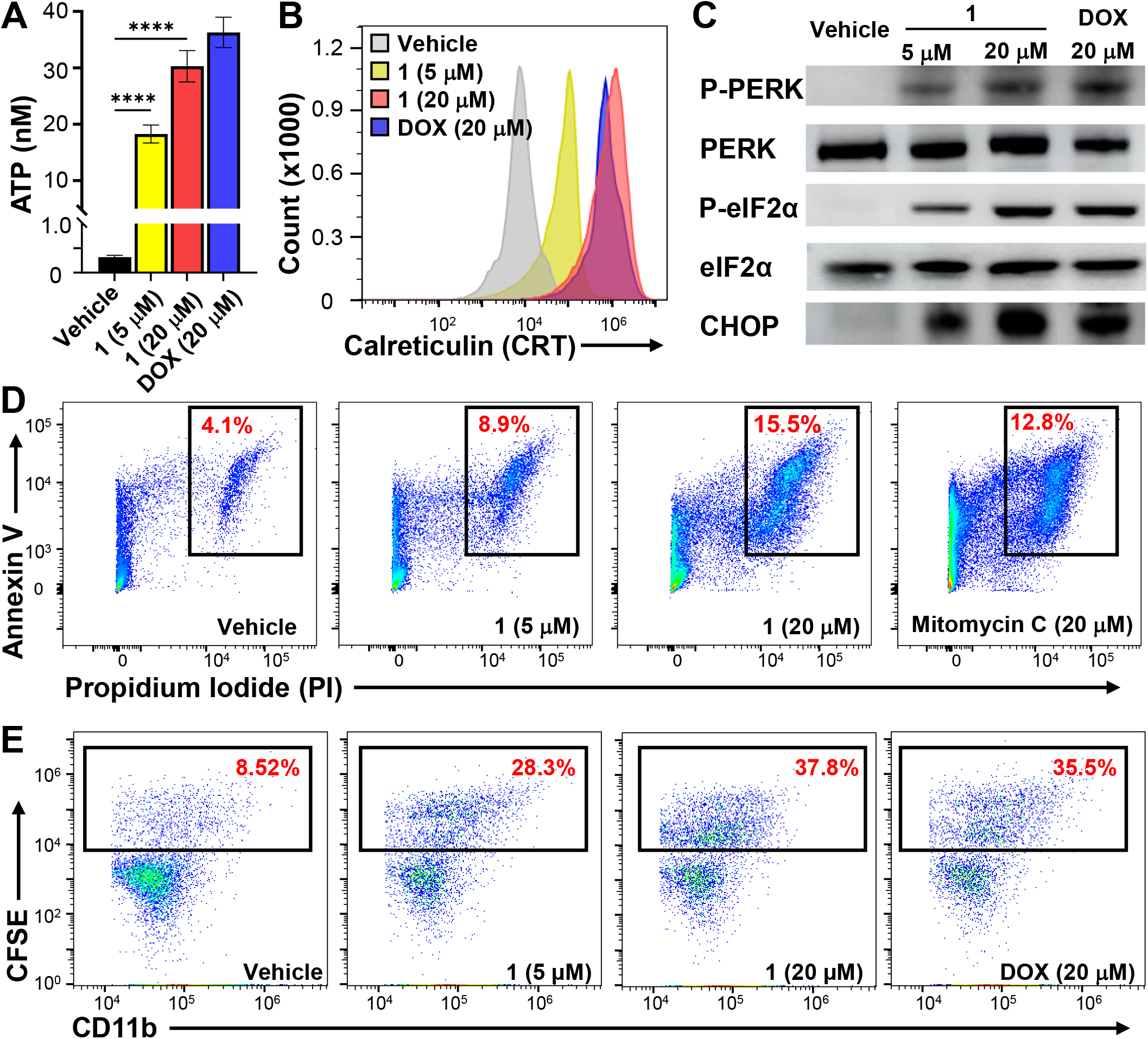
Aspercyclicin A (1) induced immunogenic cell death in PMP501-1 cells. (**A**) The extracellular ATP increased in cells treated with 5 µM (yellow) or 20 µM (red) of compound **1** compared to the DMSO (vehicle control, black). Doxorubicin (DOX, 20 µM) was used as positive control (blue). Data were acquired from three independent experiments (\*\**p* < 0.01). (**B**) An increase in CRT exposure was observed after treatment with 5 µM compound **1** (yellow), 20 µM compound 1 (red), or 20 µM DOX (positive control) (blue), compared to DMSO (vehicle control, gray) treatment. (**C**) Western blotting analysis of ER stress-related signaling in cells treated with 5 or 20 µM of compound **1** for 24h, showing the increased expression of P-PERK, P-eIF2α, and CHOP. DOX (20 µM) was used as positive control. (**D**) Representative flow cytometry plots using Annexin V-FITC/PI staining for apoptosis. Cells were treated with 5 µM or 20 µM of compound **1** for 24h. The percentage of cells with both Annexin V and PI positive (number in red at the top right), relative to the total cell population, showing potential late apoptosis. Mitomycin C (20 µM) was used as positive control. (**E**) Phagocytosis of compound **1**-treated cells by THP-1-derived macrophages. The treated cancer cells were stained by CFSE, and the THP-1-derived macrophages were stained by CD11b antibody. After six hours of co-culture, cells were analyzed by flow cytometry for the co-occurrence of the CFSE and CD11b signals. The percentage of cells with both CFSE and CD11b positive (number in red at the top right), relative to the total THP1 cell population, increased significantly in the treatment with 5 µM and 20 µM of compound **1**. DOX (20 µM) was used as positive control. For all of the above experiments, the DMSO-treated cells served as the vehicle control.

Finally, we explored more on ICD-mediated cancer cell death of aspercyclicins. We performed propidium iodide (PI) and fluorescein isothiocyanate (FITC)-conjugated Annexin V (Annexin V-FITC) staining assay. The staining results indicated the occurrence of late apoptosis in PMP501-1 cells following a 24-hour treatment with **1** (Figure 7D). Interestingly, the morphological alterations observed in PMP501-1 cells following treatment with **1** diverged from the typical features associated with late-stage apoptosis, thereby suggesting an unconventional apoptosis mechanism which supported ICD-mediated cell death. It is known that ICD dying cancer cells are able to induce antigen presenting cell (APC) maturation and to improve the ability to be engulfed by APC through phagocytosis^51^. Therefore, we tested if compound **1** treatment enhances phagocytosis of cancer cells by human macrophages. Specifically, we performed phagocytosis assay by directly culturing compound **1**-treated PMP501-1 cancer cells with THP-1-derived macrophages, followed by analyzing the cells with flow cytometry. As shown in Figure 7E, the ratio of phagocytosis of cancer cells by macrophages doubled when cancer cells were treated with **1** compared to the vehicle. Overall, these findings indicate that **1** effectively induces immunogenic death of various cancer cells, leading to their downstream phagocytic clearance by macrophages without toxicity to normal cells. Thus, aspercyclicins represent a novel class of broad-spectrum cancer immunogenic chemotherapeutic leads.

## Discussion

Recently, ICD has garnered significant attention due to its unique ability to kill cancer cells through induction of a robust and specific immune response ^52^. Discovering chemical ICD inducers holds promises for development of immunogenic chemotherapy that combines the advantages of chemotherapy and immunotherapy. Although promising, only a few small molecule ICD inducers have been discovered. Natural products are one of the most important sources for discovery of ICD inducers. The discovery of aspercyclicins contributed a novel class of compounds to the limited number of existing ICD-inducing small molecules. We identified a potential mechanism of action of aspercyclicins, revealing their ability to trigger ICD against cancer cells and potentially induce protective antitumor immunity. Importantly, aspercyclicins exhibited broad-spectrum activity in inducing ICD among multiple cancer cell lines, including a lung cancer cell line LLC. This observation is particularly encouraging in terms of that lung cancer patients often fail to respond to immunotherapy effectively^53,54^. Notably, the activity of aspercyclicins targeted PMP, a rare but severe cancer that currently lacks effective chemotherapy. Previously, no screening platform has been developed for PMP, limiting ex vivo studies and effective drug discovery for this cancer. Our work to establish stable, patient-derived PMP cell lines with favorable morphology for screening, provides a significant advancement in PMP anti-cancer drug discovery. Our platform facilitated rapid prioritization and identification of a new class of anticancer leads, the aspercyclicins. While capable of inducing ICD effects in various types of cancer cells, remarkably, the aspercyclicins showed minimal cytotoxicity against normal cells, further strengthening their potential effectiveness in treating cancers without common adverse reactions as observed for traditional cancer chemotherapeutics.

Recent microbial natural product discovery efforts have revealed that a substantial portion of natural products could result from unexpected processes, including enzymatic catalysis encoded by unclustered genes and nonenzymatic chemistries^39,55–59^. Novel metabolites produced from such processes cannot be predicted by currently existing bioinformatics tools and could be easily missed by bottom-up genome mining approaches^55^. In contrast, our forward chemical genetic screening, equipped with advanced cheminformatics tools, directly prioritized a class of hits that possessed both complex structures and significant bioactivity, leading to the discovery of aspercyclicin family. The chimeric nature of aspercyclicins not only adds to cryptic chemical coupling phenomenon but also signifies that microbes could evolve to utilize different biosynthetic pathways, transforming their products into chimeras with more diverse biological functions. Importantly, decoding these nature-inspired chimeric constructions would enhance the diversity of potential pharmacophores and inspire both chemical and biological synthesis, thereby developing drug leads more efficiently.

The push to uncover novel chemistry in natural products positions the discovery and elucidation of complex molecules as a promising frontier in drug discovery. However, the unique characteristics of these molecules can also present challenges for their isolation and structure elucidation. Aspercyclicins, for example, exhibit several unideal qualities that make their study complicated: i). The 1,4-dihydro-1,4-epoxynaphthalene motif together with the geodin moiety is proton-deficient, featuring a 21-carbon skeleton with up to 17 contiguous quaternary carbons and challenging NMR-based structural elucidation; and ii). Although potent, aspercyclicins are minor components of the extract and would be likely overlooked by direct isolation. As such, we used both bioactivity-based molecular networking and MicroED analysis to expedite our prioritization, discovery, and structural elucidation. Our utilization of integrative technology is crucial in achieving discovery and unambiguous structural determination of this class of minor yet active natural products. These results also guided subsequent MS-based metabolomics, contributing to the discovery of aspercyclicin analogs. Previously, innovative pipelines have been developed towards automated, large-scale, high-capacity, and high throughput methods for fractionation of natural product extracts, such as the NCI NPNPD program^60,61^. These sample processing methods can rapidly yield a sufficient amount of subfractions for both cell-free and cell-based bioassays (e.g., the NCI-60 activity testing) and also introduce spectroscopic and spectrometric analyses for dereplication of known compounds. Our discovery pipeline combines historically validated methods with modern techniques, taking advantage of cheminformatics and structure determination advances to introduce knowledge of bioactive, novel structural class early in the isolation workflow and expedite the discovery and structure elucidation of structurally novel and functionally useful molecules. Thus, such advancements have enabled our discovery of a remarkable array of natural products that were previously missed. The streamlined and informatics-based discovery platform reported here is widely applicable to diverse settings and is envisioned to reveal more minor yet active compounds, thereby expanding the scope of potential small molecule drug leads.

In sum, the discovery and characterization of aspercyclicins exemplify the benefits of simultaneously prioritizing novel chemistry and recognizing minor yet active components in natural product-based drug discovery. Limitations in identifying such compounds due to the technological constraints are now being overcome by new and refined methodologies as exemplified by our discovery platform reported here. The novel skeleton and broad-spectrum ICD-inducing activity of aspercyclicins, combined with their low toxicity to normal cells, may inspire further efforts to develop them into a new immunogenic chemotherapeutic drug.

## Acknowledgments

This work is partially funded by the NIH grant 1R35GM150565 and the National Organization for Rare Disorders (NORD) ACPMP pilot grant. TSB acknowledges funding from NIH U19AI142720. We thank Miss Mary Caitlin King of Mercy Medical Center for help with acquiring patient PMP samples, Dr. Perry J. Pellechia and Miss Toni Johnson from University of South Carolina (USC) NMR Facility for help with acquiring NMR data, Drs. Michael D. Walla and William E. Cotham from USC Mass Spectrometry Facility for acquiring HRMS data, Dr. Mark D. Smith from USC X-Ray Diffraction Facility for acquiring X-ray crystallography data, Drs. Gregory Morrison and Hans-Conrad zur Loye from USC Department of Chemistry and Biochemistry for acquiring PXRD data, Dr. Ying Li, a visiting scholar at Department of Chemistry and Biochemistry USC. XtalPi and Drs. David A. Delgadillo, Jessica E. Burch, Yanping Qiu, and Hosea M. Nelson at California Institute of Technology for assistance in MicroED data acquiring and interpretation, as well as Dr. Pengpeng Zhang at Yale University and Dr. Jia-Xuan Yan at Merck & Co., Inc. for helpful discussions.

## Associated Content

### Supporting Information

Full experimental details, NMR data and structure elucidation of all isolated compounds, crystal structure information, bioactivity-based molecular networking details, and bioactivity data of all extracts and fractions. (PDF)

### Supplementary File

Information of 100 strains, including taxonomic details, source, 16S or ITS sequences. (Excel)

### Author Contributions

D.X., H.C., D.F., T.S.B., T.L.T., Q.W. and J.L. designed research; D.X., M.X., and Q.W. isolated and characterized the compounds; D.X., M.D.M., X.L., E.A.O., C.P., and Z.S prepared the strain crude extract library; D.X., Y.H., G.G., M.K.R., Y.W., Y.L., H.C., D.F., and T.L.T. developed and carried out the cell assays; A.S. provided patient samples for PMP cell development; Q.W., D.X., T.S.B., and J.L. conducted the metabolomics analysis; Q.W., D.X., and J.L. wrote the manuscript, and T.S.B. revised it; All authors have reviewed, revised, and approved the final version of the manuscript.

